# Refinement of anesthetic choice in procedures preceding psychopharmacological studies

**DOI:** 10.1101/390583

**Authors:** LS Herbst, T Gaigher, AA Siqueira, SRL Joca, KN Sampaio, V Beijamini

## Abstract

**Rationale:** Previous studies indicated that some general anesthetics induce long-term antidepressant and/or anxiolytic-like effects. This raises the concern about the use of anesthesia in surgeries that precede psycopharmacological tests, since it may be a potential bias on results depending on the experimental design used.

**Objectives:** To evaluate whether commonly used general anesthetics in surgeries preceding psychopharmacological tests would affect rat behavior in tests predictive of antidepressant or anxiolytic-like effects. We also evaluated whether prior anesthesia would interfere in the detection of the antidepressant-like effect of imipramine or the anxiolytic-like effect of diazepam.

**Methods:** We tested if a single exposure to subanesthetic or anesthetic doses of 2,2,2-tribromoethanol, chloral hydrate, thiopental or isoflurane would change rat’s behavior in the forced swimming test (FST) or in the elevated plus-maze (EPM) test, at 2 hours or 7 days after administration.

**Results:** Previous anesthesia with the aforementioned anesthetics did not change rat behavior in FST per se nor it changed the antidepressant-like effect induced by imipramine treatment. Rats previously anesthetized with tribromoethanol or chloral hydrate exhibited, respectively, anxiogenic-like or anxiolytic-like behavior in the EPM. Prior anesthesia with thiopental or isoflurane did not produce any *per se* effect in rat behavior in the EPM nor disturbed the anxiolytic-like effect of diazepam.

**Conclusion:** Our results suggest that, in our experimental conditions, tribromoethanol and chloral hydrate are improper anesthetics for surgeries that precede behavioral tests related to anxiety. Isoflurane or thiopental may be suitable for anesthesia before evaluation in animal models predictive of antidepressant or anxiolytic-like effect.

## 1 Introduction

Rodent models are frequently used for understanding the pathophysiology of various diseases and for developing new therapeutic strategies (Flecknell 1988; Stokes et al. 2009; Gargiulo et al. 2012). Procedures such as stereotactic surgery for implantation of cannulas/probes or electrodes in the central nervous system (CNS) are fundamental tools for many applications, such as targeted drug delivery, injection of anatomical tracers, site-specific brain lesions, microdialysis studies, among others (Yoshida et al. 2015). Given that stereotaxic surgery is an invasive and painful procedure, the administration of anesthetics (general and/or local) is mandatory.

General anesthesia in laboratory animals consists in loss of consciousness, analgesia, muscle relaxation and suppression of reflexes (Flecknell 2009). An ideal anesthetic should promote fast and reversible narcosis with as little pain as possible, cause few side effects, including reduced post-anesthetic impact on the CNS, be safe for the manipulators and still be easy to administer (Gargiulo et al. 2012; Hüske et al. 2016). Thus, the depth of anesthesia and the potential adverse effects of the anesthetic, such as hypothermia, cardiovascular or respiratory depression, should be taken into account when choosing the most appropriated anesthetic (Arras et al. 2001; Buitrago et al. 2008; Hüske et al. 2016). Beyond the safety and effectiveness of the chosen anesthetic, its potential bias on experimental variables should also be taken into consideration.

General anesthetics may be classified according to the route of administration as injectable or inhalation (Flecknell 2009). Injectable anesthetics are frequently used in laboratory rodent surgeries due to their low cost and convenient application (Arras et al. 2001; Buitrago et al. 2008; Stokes et al. 2009; Hüske et al. 2016). However, injectable anesthesia is quite a challenge regarding the choice of dose to ensure adequate depth of anesthesia and, at the same time, avoid marked changes in cardiovascular and respiratory functions (Smith 1993; Janssen et al. 2004; Gargiulo et al. 2012). Inhalation anesthetics allow easy maintenance of the surgical anesthetic plane, providing greater safety than the injectable ones (Roth et al. 2002; Zuurbier et al. 2002; Janssen et al. 2004; Gargiulo et al. 2012). Also, inhalation agents are eliminated through respiratory route, usually enabling a rapid recovery of anesthesia. On the other hand, given that they require a device for appropriately delivery of the anesthetic, the total cost of anesthesia may be higher (Buitrago et al. 2008; Cesarovic et al. 2010). Further, inhalation anesthetics may induce respiratory depression and sometimes require intubation procedure (Gargiulo et al. 2012; Tsukamoto et al. 2015).

Stokes and coworkers (2009) showed that ketamine, chloral hydrate, pentobarbital and isoflurane are typical anesthetics in rodent surgical procedures. However, the use o pentobarbital is declining because of its decreasing availability (Richardson and Flecknell 2005; Priest and Geisbuhler 2015). Accordingly, thiopental may be a barbiturate option in short-time procedures (Sumitra et al. 2004; Gaertner et al. 2008). Tribromoethanol has also been a common option due to its relative safety, rapid-onset and fast recovery of anesthesia (Gaertner et al. 2008; Maheras and Gow 2013).

In recent years, a growing body of evidence has shown that anesthetics can induce long-term brain changes that may affect animal’s behavior (Colon et al. 2017). For instance, it is now well recognized that subanesthetic and anesthetic doses of the general anesthetic ketamine induce a fast and persistent antidepressant-like effect in rodents (Yilmaz et al. 2002; Maeng et al. 2008). Ketamine also showed acute anxiolytic-like effect at subanesthetic doses in the elevated plus-maze (EPM) test (Engin et al. 2009). Considering that in neuropsychopharmacological studies it is common to perform surgical procedures under anesthesia before the behavioral evaluation, a better understanding of the potential bias of general anesthetics in this kind of experimental design is imperative.

Based on that, we addressed whether the following general anesthetics commonly used in rodents during surgical procedures, tribromoethanol, chloral hydrate, thiopental or isoflurane, would affect rat behavior in tests predictive of antidepressant or anxiolytic effects. Moreover, we assessed whether prior anesthesia would interfere with the detection of the antidepressant-like effect of imipramine or the anxiolytic-like effect of diazepam. In addition, we also evaluated the potential antidepressant or anxiolytic-like effects of subanesthetic doses of the aforementioned anesthetics.

## 2 Materials and methods

### 2.1 Animals

Adult (8-10 weeks old) and naive male Wistar rats (280-350 g) were obtained from the breeding colony of Federal University of Espirito Santo and allowed to acclimatize in our laboratory for 4 weeks before intervention start. Rats were kept in groups of five animals per cage (49 x 34 x 26 cm) in a temperature-controlled room (24 ± 2◦C) under a 12 h/12 h light/dark cycle (light on at 6:30 a.m.) with food and water available ad libitum. Rats were fed a standard chow diet (Nuvilab CR1, Quintia S.A. Brazil). Cages were changed once a week. The total number of rats used was 543. The experimental procedures were approved by the local Committee for the Ethical Use of Animals in scientific research (CEUA-UFES, 58/2015).

### 2.2 Drugs and Treatment

The injectable general anesthetics were freshly diluted in saline and administered by the intraperitoneal (i.p.) route at 3 different doses (2 subanesthetic and 1 anesthetic doses), as named: 2,2,2-tribromoethanol (Sigma-Aldrich, MO, USA) 40, 90 and 250 mg/Kg; chloral hydrate (VETEC, SP, Brazil) 50, 150 and 400 mg/kg; thiopental (Cristália, SP, Brazil) 5, 15 and 40 mg/kg. For each anesthetic, the highest dose is the anesthetic one, according to previous studies from literature (Vachon et al. 2000; Sumitra et al. 2004; Ajadi et al. 2013).

Isoflurane (Cristália, SP, Brazil) was delivered by an inhalation anesthesia apparatus (Bonther®, SP, Brazil), which consists of a vaporizer coupled to a digital flowmeter and to a translucent plastic chamber. As a carrier gas, pressurized air was used at flow rate of 2l/min. The animals were placed in the chamber for induction and maintenance of anesthesia. The subanesthetic concentrations used were 0.5 and 1.5%. Anesthesia was inducted with 4% of isoflurane (approximately 5 minutes for induction) and maintained with 2% (Maud et al. 2014). Administration was performed in a room with temperature varying between 22 and 24 ºC.

Anesthesia was confirmed by 2 noxious stimuli: pinching of the tail (tail pinch reflex) and interdigital webbing (pedal withdrawal reflex).

Imipramine 15mg/kg (Sigma-Aldrich, MO, USA) and diazepam 2.5mg/kg (Cristália, SP, Brazil) were used as positive controls for antidepressant (Porsolt et al. 1978) or anxiolytic-like effect (Pellow et al. 1985), respectively.

### 2.3 Behavioral tests

#### 2.3.1 Forced Swimming Test (FST)

The FST was performed The test was performed in 2 swimming sessions using a cylinder (23 cm diameter and 52 cm height) containing 30 cm of water with temperature between 22 e 24ºC. In the pre-test session, rats were placed on the cylinder to swim for 15 minutes. Twenty-four hours later, the animals were tested for 5 minutes (test session). FST behavior was videorecorded and analyzed by a researcher who was blind to the treatment groups. The frequency of climbing, swimming and immobility was scored every 5 seconds, according to the procedure described by Detke and colleagues (1995). Climbing consists in vigorous vertically movements with forepaws against the wall of the cylinder. Swimming is identified as horizontal movements across the water surface. Immobility consists in minor movements strictly necessary to maintain the animal’s head above water.

#### 2.3.2 Elevated Plus-Maze (EPM) Test

The EPM apparatus is a plus-shaped maze formed by 2 enclosed arms (surrounded by walls of 40cm high) and 2 open arms. The open arms have a transparent border of 1cm in height to prevent animal’s fall. In the EPM test, each animal was placed in the center of the platform and allowed to freely explore the maze for 5 min. An experimenter blind to drug treatment analyzed the recorded sections to compute anxiety-like behaviors: percentage of open arms entries and percentage of time spent in open arms. The number of enclosed arms entries was also evaluated as an index of exploratory activity (Pellow et al. 1985).

#### 2.3.3 Open Field Test (OF)

The apparatus consists in a square box (1m^2^) with black floor and surrounded by walls of 30cm of weight. In the test session, each rat was placed in center of the OF and allowed to freely explore the apparatus for 5 minutes. The test sessions were videorecorded and analyzed by the ANY-maze™ 4.98 software (Stoelting, Wood Dale, IL, USA) to obtain the total distance traveled (in meters), as an index of locomotor activity (Prut and Belzung 2003).

### 2.4 Experimental groups and procedures

All behavioral procedures were performed between 1 and 6 p.m. by the same experimenter. Rats were habituated to the behavioral rooms 3 hours before testing and were randomly assigned for one of the experiments described bellow. The first sets of experiments (Fig. 1) were designed to evaluate acute (2 hours after administration) and/or long-term effects (7 days after administration) of the following general anesthetics: tribromoethanol, chloral hydrate, thiopental and isoflurane. We chose 2 hours of interval between administration and the first test based on a pilot study in which we observed full anesthesia recovery of the animals after this period with all anesthetics tested. We defined 7 days of interval to evaluate long-term effects of the anesthetics because this interval is usually assumed as an adequate recovery interval between stereotaxic surgery and behavioral tests.

**Fig. 1.**
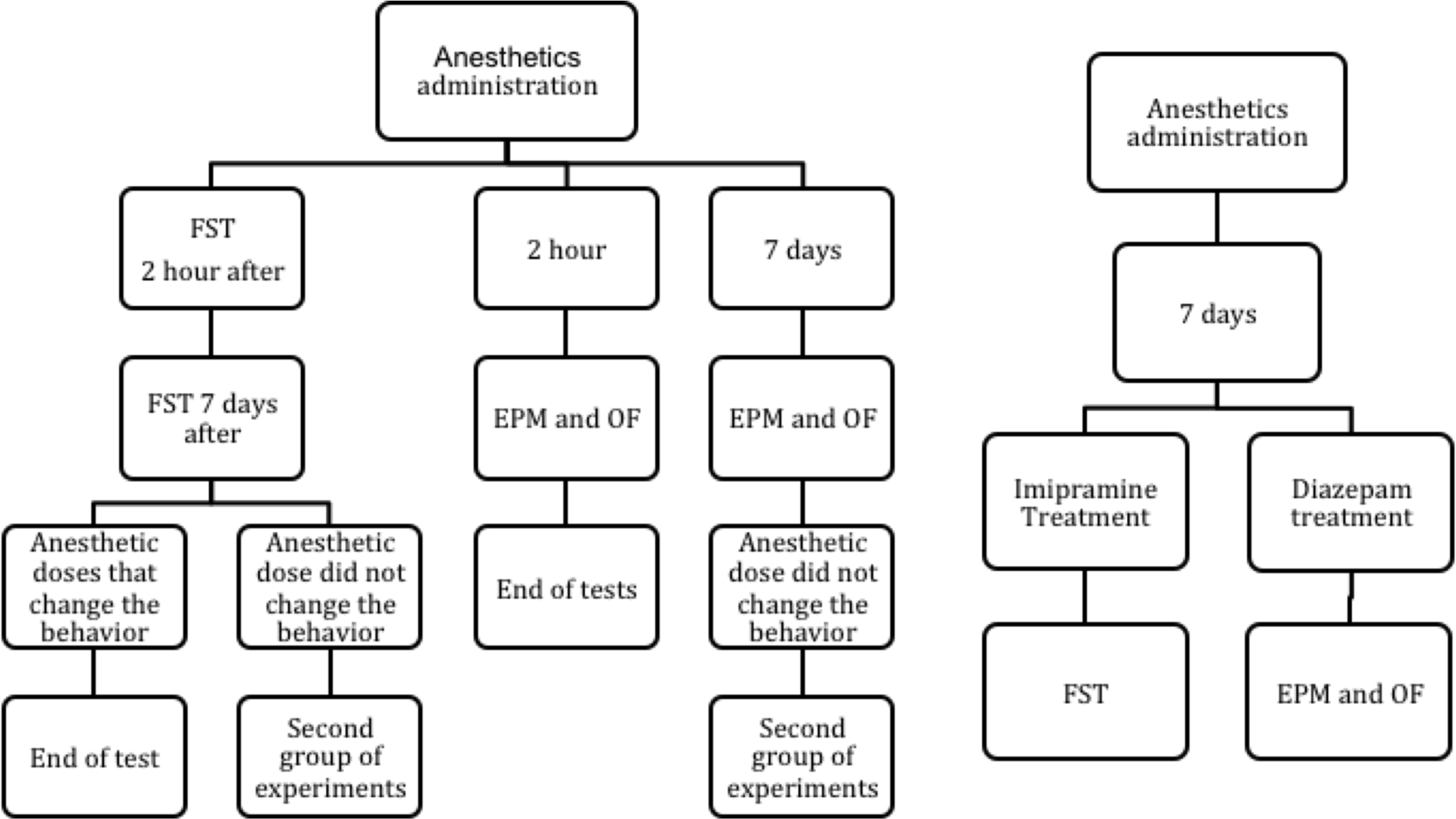
Experimental design: Left panel represents the first group of experiments and right panel represents the second group of experiments. FST (Forced swimming test); EPM (elevated plus-maze); OF (open field).

We defined the experimental design respecting the ethical recommendation of 3R’s and the methodological limitations of the behavioral tests. In this line, the same rats were tested 2 hours and 7 days after anesthetic administration in the FST (Mezadri et al. 2011). Imipramine was used as a positive control of antidepressant-like effect. In EPM tests, 2 different groups of animals were used: one group was tested 2 hours after anesthetic administration and another independent group was tested 7 days after administration of anesthetics to avoid one trial tolerance phenomenon previously described (File and Zangrossi 1993). Diazepam was used as a positive control of anxiolytic-like effect in the acute tests. Thirty seconds after the EPM test, the same animals were evaluated in the OF. We also tested a reduced number of positive controls in each experiment. Thus, for statistical analysis, we created a pool of positive controls used in all experiments. The experimental groups were detailed in the **table S1** of the supplementary material.

The second set of experiments was planned to investigated if previous administration of an anesthetic would affect the detection of the antidepressant-like effect induced by imipramine in the FST or the anxiolytic-like effect induced by diazepam in the EPM test **(Fig. 1).** We chose to evaluate only the general anesthetics that did not modify, *per se*, the animal behavior in the FST or in the EPM 7 days after anesthetic administration. Thus, rats received an anesthetic dose of tribromoethanol, chloral hydrate, thiopental or saline and, 7 days later, they received 1 injection imipramine (15mg/kg i.p.) or saline and were tested in the FST 1 hour after the last injection. Another group of animals were exposed to isoflurane or air and, 7 days later, they treated with imipramine or saline, 1 hour before the FST. The same protocol were employed to test whether thiopental or isoflurane affect the anxiolytic-like effect of diazepam (2.5mg/kg, i.p.) in the EPM test. The experimental groups were detailed in the **table S2** of the supplementary data.

### 2.5 Statistical analysis

Data were represented as mean ± standard error of mean (S.E.M). All analyses were performed through the SPSS for Windows© (version 2.0). Levene’s test was performed to evaluate homoscedasticity. In cases of absence of homoscedasticity, the data were square root-transformed and analyzed. Data from FST with 2 test sessions were analyzed by two-way Analysis of Variance (ANOVA) with repeated measures and *post hoc* of Duncan when appropriated. Data from EPM and OF were analyzed by one-way ANOVA followed by Duncan’s test. The results from positive controls imipramine and diazepam were compared with those of saline groups using the Student’s t test for independent measures. Data from the second battery of tests were analyzed by two-way ANOVA and *post hoc* of Duncan when appropriated. The significance was setup as p<0.05.

## 3 Results

Table 1 shows the effects of subanesthetic and anesthetic doses of tribromoethanol, chloral hydrate, thiopental and isoflurane in rats submitted to the FST. None of tribromoethanol doses interfered with climbing, swimming or immobility frequencies in the FST, either 2 hours or 7 days after anesthetic administration. Chloral hydrate 150mg/kg decreased the immobility frequency acutely and 7 days after administration, compared to saline group, while the other doses did not change behavioral parameters in the FST. The anesthetic dose of thiopental (40mg/kg) decreased immobility and increased swimming frequency acutely. The other tested doses did not impact FST behaviors acutely or 7 days later. Isoflurane did not affect any behavior in the FST, 2 h or 7 days after exposure. The positive control imipramine increased climbing and decreased immobility frequency without interfering in swimming behavior. The statistical data were detailed in **table 1**.

**Table 1.** Effects of a single exposure to tribromoethanol, chloral hydrate, thiopental or isoflurane in rats submitted to the forced swimming test 2 h and 7 days after anesthetic administration.

| FORCED SWIMMING TEST |  |  |  |  |
| --- | --- | --- | --- | --- |
| Treatment (n) |  | Climbing | Swimming | Immobility |
| Saline (8) | Test | 19.37 ± 3.34 | 3.25 ± 0.75 | 35.87 ± 4.48 |
|  | Retest | 14.16 ± 5.16 | 4.16 ± 1.49 | 41.33 ± 4.96 |
| Tribromoethanol 40mg/kg (7) | Test | 32.14 ± 5.23 | 4.14 ± 0.91 | 21.85 ± 5.02 |
|  | Retest | 25.71 ± 7.44 | 5.00 ± 2.14 | 27.14 ± 7.06 |
| Tribromoethanol 90mg/kg (9) | Test | 20.44 ± 6.28 | 5.88 ± 1.74 | 29.88 ± 5.10 |
|  | Retest | 10.55 ± 2.09 | 3.11 ± 0.69 | 37.44 ± 5.95 |
| Tribromoethanol 250mg/kg (8) | Test | 19.00 ± 4.14 | 4.00 ± 0.90 | 35.12 ± 4.91 |
|  | Retest | 20.33 ± 4.92 | 6.22 ± 1.25 | 32.44 ± 5.79 |
| Imipramine<sup>a</sup> (8) | Test | 42.75 ± 5.57<sup>#</sup> | 6.00 ± 2.82 | 9.12 ± 3.63<sup>#</sup> |
|  | Retest | 19.14 ± 1.80 | 3.28 ± 0.94 | 32.42 ± 1.32 |
| Session |  | [F<sub>(1,28)</sub>=10.067;p<0.004] | [F<sub>(1,28)</sub>=0.016;p=0.900] | [F<sub>(1,28)</sub>=2.276;p=0.114] |
| Treatment |  | [F<sub>(3,28)</sub>=1.553;p=0.225] | [F<sub>(3,28)</sub>=0.148;p=0.930] | [F<sub>(3,28)</sub>=1.381;p=0.271] |
| Interaction |  | [F<sub>(3,28)</sub>=1.052;p=0.386] | [F<sub>(3,28)</sub>=1.483;p=0.242] | [F<sub>(3,28)</sub>=0.300;p=0.825] |
| Treatment (n) |  | Climbing | Swimming | Immobility |
| Saline (15) | Test | 21.33 ± 2.48 | 2.86 ± 0.66 | 32.00 ± 3.01 |
|  | Retest | 22.60 ± 4.20 | 1.93 ± 0.35 | 33.73 ± 4.15 |
| Chloral hydrate 50mg/kg (10) | Test | 20.40 ± 5.29 | 3.70 ± 0.89 | 31.90 ± 5.29 |
|  | Retest | 20.50 ± 6.68 | 2.60 ± 0.54 | 35.60 ± 6.94 |
| Chloral hydrate 150mg/kg (12) | Test | 31.50 ± 4.04 | 3.25 ± 1.05 | 18.91 ± 5.00* |
|  | Retest | 33.58 ± 5.36 | 3.00 ± 0.57 | 19.75 ± 5.31* |
| Chloral hydrate 400mg/kg (9) | Test | 23.88 ± 2.19 | 1.77 ± 0.49 | 27.44 ± 2.58 |
|  | Retest | 24.88 ± 3.23 | 2.88 ± 1.24 | 27.88 ± 2.38* |
| Imipramine<sup>a</sup> (8) | Test | 42.75 ± 5.57<sup>#</sup> | 6.00 ± 2.82 | 9.12 ± 3.63<sup>#</sup> |
|  | Retest | 19.14 ± 1.80 | 3.28 ± 0.94 | 32.42 ± 1.32 |
| Session |  | [F<sub>(1,42)</sub>=27.419;p=0.471] | [F<sub>(1,42)</sub>=1.903;p=0.565] | [F<sub>(1,42)</sub>=0.333;p=0.567] |
| Treatment |  | [F<sub>(3,42)</sub>=1.755;p=0.170] | [F<sub>(3,42)</sub>=4.455;p=0.604] | [F<sub>(3,42)</sub>=3.474;p=0.024] |
| Interaction |  | [F<sub>(3,42)</sub>=3.643;p=0.975] | [F<sub>(3,42)</sub>=4.997;p=0.457] | [F<sub>(3,42)</sub>=0.043;p=0.988] |
| Treatment (n) |  | Climbing | Swimming | Immobility |
| Saline (8) | Test | 26.62 ± 5.79 | 1.00 ± 0.37 | 29.87 ± 5.59 |
|  | Retest | 36.62 ± 5.95 | 1.62 ± 1.10 | 19.87 ± 6.11 |
| Thiopental 05mg/kg (7) | Test | 20.71 ± 3.06 | 3.57 ± 2.59 | 34.14 ± 4.67 |
|  | Retest | 31.42 ± 5.87 | 1.00 ± 0.69 | 25.57 ± 5.84 |
| Thiopental 15mg/kg (7) | Test | 22.57 ± 11.85 | 1.57 ± 0.75 | 34.57 ± 4.03 |
|  | Retest | 25.14 ± 5.20 | 1.00 ± 0.37 | 31.71 ± 5.51 |
| Thiopental 40mg/kg (7) | Test | 29.71 ± 17.37 | 15.85 ± 3.94* | 11.00 ± 3.57* |
|  | Retest | 32.14 ± 4.07 | 2.00 ± 0.87 | 23.57 ± 4.82 |
| Imipramine<sup>a</sup> (8) | Test | 42.75 ± 5.57 | 6.00 ± 2.82 | 9.12 ± 3.63<sup>#</sup> |
|  | Retest | 19.14 ± 1.80 | 3.28 ± 0.94 | 32.42 ± 1.32 |
| Session |  | [F<sub>(1,25)</sub>=8.658;p=0.007] | [F<sub>(1,25)</sub>=11.988;p=0.002] | [F<sub>(1,25)</sub>=1.057;p=0.314] |
| Treatment |  | [F<sub>(3,25)</sub>=0.608;p=0.616] | [F<sub>(3,25)</sub>=8.143;p=0.001] | [F<sub>(3,25)</sub>=2.086;p=0.128] |
| Interaction |  | [F<sub>(3,25)</sub>=1.078;p=0.376] | [F<sub>(3,25)</sub>=7.818;p=0.001] | [F<sub>(3,25)</sub>=5.715;p=0.004] |
| Treatment (n) |  | Climbing | Swimming | Immobility |
| Air (5) | Test | 25.40 ± 7.54 | 2.20 ± 0.73 | 29.60 ± 8.49 |
|  | Retest | 23.16 ± 6.23 | 1.16 ± 0.86 | 34.50 ± 6.65 |
| Isoflurane 0,5% (7) | Test | 37.57 ± 5.93 | 0.00 ± 0.00 | 21.28 ± 5.84 |
|  | Retest | 26.85 ± 8.04 | 0.00 ± 0.00 | 30.57 ± 7.78 |
| Isoflurane 1,5% (7) | Test | 25.14 ± 6.04 | 2.28 ± 0.91 | 31.57 ± 5.68 |
|  | Retest | 20.14 ± 4.78 | 1.57 ± 0.68 | 34.00 ± 5.00 |
| Isoflurane 4,0% (7) | Test | 22.57 ± 2.71 | 2.14 ± 0.76 | 33.42 ± 2.45 |
|  | Retest | 20.00 ± 2.00 | 0.71 ± 0.35 | 38.14 ± 2.06 |
| Imipramine<sup>a</sup> (8) | Test | 42.75 ± 5.57 | 6.00 ± 2.82 | 9.12 ± 3.63<sup>#</sup> |
|  | Retest | 19.14 ± 1.80 | 3.28 ± 0.94 | 32.42 ± 1.32 |
| Session |  | [F<sub>(1,22)</sub>=2.670;p=0.116] | [F<sub>(1,22)</sub>=4.099;p=0.055] | [F<sub>(1,22)</sub>=3.481;p=0.075] |
| Treatment |  | [F<sub>(3,22)</sub>=1.021;p=0.403] | [F<sub>(3,22)</sub>=2.972;p=0.054] | [F<sub>(3,22)</sub>=0.720;p=0.550] |
| Interaction |  | [F<sub>(3,22)</sub>=0.660;p=0.585] | [F<sub>(3,22)</sub>=0.711;p=0.556] | [F<sub>(3,22)</sub>=0.363;p=0.781] |

<sup>a</sup> Student's t test independent measure. # $p < 0.05$ imipramine vs control group (saline or air). \* $p < 0.05$ vs saline group. ANOVA followed by Duncan test

Previous anesthesia with tribromoethanol 250mg/kg, chloral hydrate 400mg/kg, thiopental 40mg/kg or isoflurane (4% for induction and 2 % for maintenance) did not disturb the ability of imipramine in decreasing immobility and increasing climbing behaviors in the rat’s FST compared to the control group **(Fig. 2).** The statistical data were detailed in **table S3** of the supplementary material.

**Fig. 2.**
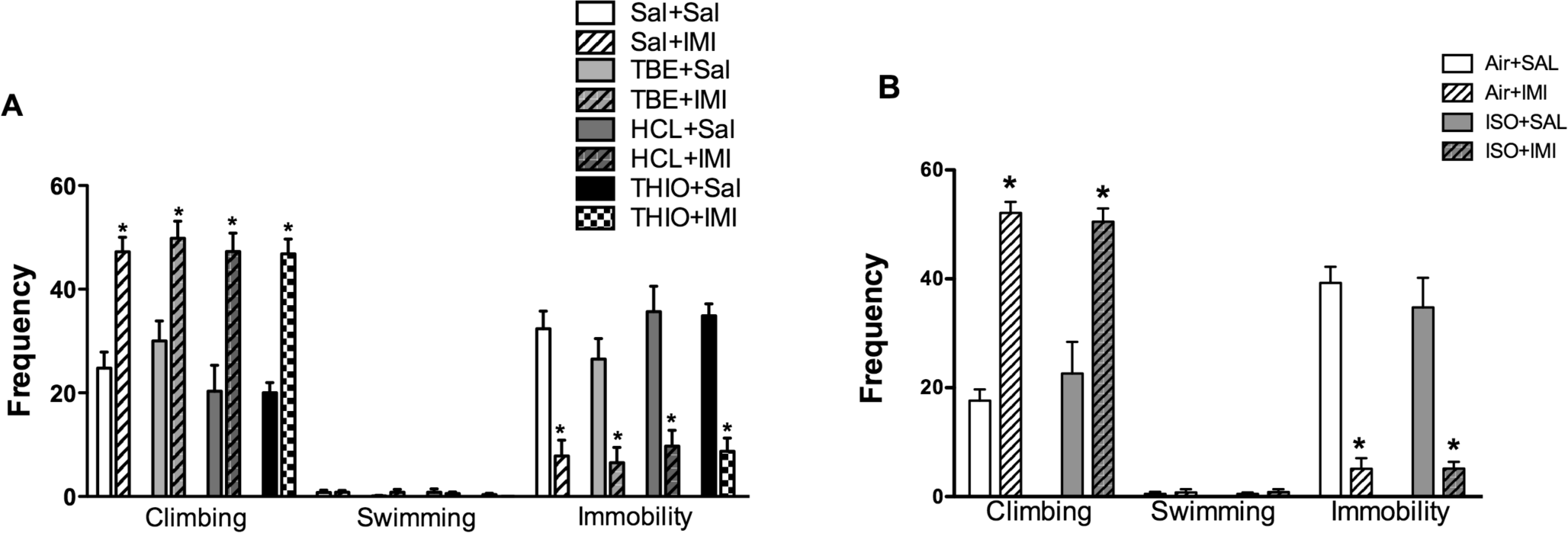
Previous anesthesia with tribromoethanol, choral hydrate, thiopental or isoflurane does not affect the antidepressant-like effect of imipramine in the rat’s forced swimming test (FST). Anesthetics were injected 7 days and imipramine 1 h before the rat’s FST. Rats treated with (A) saline-saline (SAL+SAL, n=8); saline + imipramine 15mg/kg (SAL+IMI, n=5); tribromoethanol 250mg/kg + saline (TBE+SAL, n=8); tribromoethanol + imipramine 15mg/kg (TBE+IMI, n=6), choral hydrate 400mg/kg + saline (HCL+SAL, n=6), chloral hydrate 400mg/kg + imipramine 15mg/kg (HCL+IMI, n=7) thiopental 40mg/kg + saline (THIO+SAL, n=8) and thiopental 40mg/kg + imipramine 15mg/kg (THIO+IMI, n=7). (B) air- saline (AIR-SAL, n=8); air-imipramine (AIR-IMI, n=8); isoflurane-saline (ISO-SAL, n=8); isoflurane-imipramine (ISO-IMI, n=6) *p<0,05 versus Sal-Sal group (two-way ANOVA followed by Duncan’s *post hoc* test).

As shown in **table 2**, anesthesia with tribromoethanol, 7 days before the EPM test, decreased the percentage of open arms entries but it did not affect the percentage of time spent in the open arms or the number of closed arms entries in the EPM. The subanesthetic doses of tribromoethanol did not impact rat’s behaviors in the EPM 7 days after its administration. However, tribromoethanol 90 mg/kg acute injection increased the percentage of time spent in open arms of the EPM, marginally increased the percentage of open arms entries, without changing the number of enclosed arms entries compared to saline group. The anesthetic dose of chloral hydrate (400mg/kg) enhanced the percentage of open arms entries 2 hours after its administration without changing the percentage of time spent in the open arms or the number of enclosed arms entries. Chloral hydrate 400mg/kg also raised the percentage of open arms entries and the percentage of time spent in the open arms 7 days after anesthesia but did not change the number of enclosed arms entries. Thiopental and isoflurane did not change any behavior in the EPM, 2 h or 7 days after administration. The positive control diazepam enhanced the percentage of open arms entries, the percentage of time spent in the open arms and the number of enclosed arms entries compared to the control groups.

**Table 2.** Effects of a single exposure to tribromoethanol, chloral hydrate, thiopental or isoflurane in rats submitted to the elevated plus-maze test 2 h or 7 days after anesthetic administration.

| ELEVATED PLUS MAZE |  |  |  |
| --- | --- | --- | --- |
| 2 hours |  |  |  |
| Treatment (n) | % Open Arms Entries | % Open Arms Time | Enclosed Arms Entries |
| Saline (15) | 10.83 ± 3.71 | 3.86 ± 1.66 | 5.33 ± 0.88 |
| Tribromoethanol 40mg/kg (9) | 14.19 ± 4.23 | 5.38 ± 1.58 | 5.11 ± 1.12 |
| Tribromoethanol 90mg/kg (10) | 28.71 ± 5.75 | 14.78 ± 4.53* | 500 ± 1.05 |
| Tribromoethanol 250mg/kg (9) | 21.64 ± 7.10 | 9.90 ± 3.80 | 4.33 ± 0.89 |
| Diazepam<sup>a</sup> (9) | 30.76 ± 6.69<sup>#</sup> | 21.68 ± 5.31<sup>#</sup> | 9.33 ± 1.57<sup>#</sup> |
|  | [F<sub>(3,39)</sub>=2.610;p=0.065] | [F<sub>(3,39)</sub>=2.884;p=0.048] | [F<sub>(3,39)</sub>=0.182;p=0.908] |
| 7 days |  |  |  |
| Treatment (n) | % Open Arms Entries | % Open Arms Time | Enclosed Arms Entries |
| Saline (8) | 26.70 ± 5.13 | 13.85 ± 3.57 | 26.70 ± 5.13 |
| Tribromoethanol 40mg/kg (8) | 20.74 ± 6.71 | 10.79 ± 5.18 | 20.74 ± 6.71 |
| Tribromoethanol 90mg/kg (8) | 19.55 ± 6.15 | 10.72 ± 4.23 | 19.55 ± 6.15 |
| Tribromoethanol 250mg/kg (8) | 4.91 ± 3.37* | 2.41 ± 1.64 | 4.91 ± 3.37 |
|  | [F<sub>(3,28)</sub>=3.165;p=0.040] | [F<sub>(3,28)</sub>=1.533;p=0.228] | [F<sub>(3,28)</sub>=2.268;p=0.102] |
| 2 hours |  |  |  |
| Treatment (n) | % Open Arms Entries | % Open Arms Time | Enclosed Arms Entries |
| Saline (9) | 11.11 ± 7.34 | 3.72 ± 3.03 | 2.77 ± 0.43 |
| Chloral hydrate 50mg/kg (10) | 19.32 ± 5.16 | 4.68 ± 1.49 | 4.60 ± 0.79 |
| Chloral hydrate 150mg/kg (11) | 13.55 ± 5.04 | 13.61 ± 8.67 | 5.00 ± 0.79 |
| Chloral hydrate 400mg/kg (7) | 39.61 ± 7.53* | 14.90 ± 6.22 | 4.57 ± 0.89 |
| Diazepam<sup>a</sup> (9) | 30.76 ± 6.69<sup>#</sup> | 21.68 ± 5.31 | 9.33 ± 1.57<sup>#</sup> |
|  | [F<sub>(3,33)</sub>=3.723;p=0.021] | [F<sub>(3,33)</sub>=0.934;p=0.435] | [F<sub>(3,33)</sub>=1.720;p=0.176] |
| 7 days |  |  |  |
| Treatment (n) | % Open Arms Entries | % Open Arms Time | Enclosed Arms Entries |
| Saline (7) | 6.74 ± 4.36 | 1.44 ± 1.21 | 2.71 ± 0.74 |
| Chloral hydrate 50mg/kg (8) | 12.70 ± 5.08 | 3.74 ± 2.18 | 6.00 ± 0.98 |
| Chloral hydrate 150mg/kg (6) | 7.63 ± 5.52 | 1.34 ± 0.94 | 4.83 ± 1.24 |
| Chloral hydrate 400mg/kg (5) | 34.10 ± 8.49* | 11.08 ± 1.17* | 5.40 ± 1.24 |
|  | [F<sub>(3,22)</sub>=4.213;p=0.017] | [F<sub>(3,22)</sub>=6.467;p=0.003] | [F<sub>(3,22)</sub>=2.059;p=0.135] |
| 2 hours |  |  |  |
| Treatment (n) | % Open Arms Entries | % Open Arms Time | Enclosed Arms Entries |
| Saline (10) | 16.36 ± 4.89 | 8.30 ± 2.95 | 5.50 ± 3.64 |
| Thiopental 05mg/kg (9) | 24.56 ± 7.65 | 12.97 ± 5.14 | 4.44 ± 2.01 |
| Thiopental 15mg/kg (11) | 27.64 ± 5.63 | 15.06 ± 4.26 | 5.63 ± 3.85 |
| Thiopental 40mg/kg (8) | 34.57 ± 7.28 | 19.98 ± 6.96 | 5.50 ± 2.63 |
| Diazepam<sup>a</sup> (9) | 30.76 ± 6.69 | 21.68 ± 5.31<sup>#</sup> | 9.33 ± 1.57<sup>#</sup> |
|  | [F<sub>(3,34)</sub>=1.368;p=0.269] | [F<sub>(3,34)</sub>=0.966;p=0.420] | [F<sub>(3,34)</sub>=0.330;p=0.803] |
| 7 days |  |  |  |
| Treatment (n) | % Open Arms Entries | % Open Arms Time | Enclosed Arms Entries |
| Saline (8) | 27.53 ± 7.19 | 18.88 ± 5.93 | 5.75 ± 0.83 |
| Thiopental 05mg/kg (8) | 27.22 ± 6.64 | 17.04 ± 5.24 | 6.25 ± 1.29 |
| Thiopental 15mg/kg (8) | 25.10 ± 5.59 | 10.09 ± 3.94 | 4.62 ± 0.59 |
| Thiopental 40mg/kg (8) | 14.18 ± 6.19 | 10.04 ± 5.08 | 5.62 ± 0.99 |
|  | [F<sub>(3,28)</sub>=0.962;p=0.424] | [F<sub>(3,28)</sub>=0.818;p=0.495] | [F<sub>(3,28)</sub>=0.497;p=0.687] |
| 2 hours |  |  |  |
| Treatment (n) | % Open Arms Entries | % Open Arms Time | Enclosed Arms Entries |
| Air (7) | 15.07 ± 5.69 | 4.53 ± 1.96 | 5.57 ± 1.34 |
| Isoflurane 0,5% (7) | 13.31 ± 5.03 | 6.44 ± 3.73 | 6.71 ± 0.86 |
| Isoflurane 1,5% (7) | 22.78 ± 8.61 | 6.68 ± 4.13 | 5.14 ± 0.85 |
| Isoflurane 4,0% (7) | 21.03 ± 7.95 | 5.75 ± 2.21 | 4.28 ± 0.94 |
| Diazepam<sup>a</sup> (9) | 30.76 ± 6.69 | 21.68 ± 5.31<sup>#</sup> | 9.33 ± 1.57<sup>#</sup> |
|  | [F<sub>(3,24)</sub>=0.349;p=0.790] | [F<sub>(3,24)</sub>=0.094;p=0.963] | [F<sub>(3,24)</sub>=0.977;p=0.420] |

| 7 days |  |  |  |
| --- | --- | --- | --- |
| Treatment (n) | % Open Arms Entries | % Open Arms Time | Enclosed Arms Entries |
| Air (7) | 23.81 ± 6.62 | 12.41 ± 6.14 | 4.14 ± 0.55 |
| Isoflurane 0,5% (6) | 22.22 ± 7.63 | 11.58 ± 3.77 | 6.33 ± 0.84 |
| Isoflurane 1,5% (7) | 14.68 ± 6.37 | 8.15 ± 3.03 | 6.28 ± 1.26 |
| Isoflurane 4,0% (7) | 30.39 ± 8.92 | 8.21 ± 2.60 | 4.14 ± 0.73 |
|  | [F<sub>(3,23)</sub>=0.775;p=0.520] | [F<sub>(3,23)</sub>=0.292;p=0.831] | [F<sub>(3,23)</sub>=1.946;p=0.150] |

<sup>a</sup> Student's t test independent measure. # p<0.05 diazepam vs control group (saline or air).
\*p<0.05 versus saline group. ANOVA followed by Duncan test

As prior anesthesia with thiopental or isoflurane did not change rats behaviors in the EPM *per se*, we evaluated whether anesthesia with these drugs would disturb the detection of the anxiolytic-like effect of diazepam. Previous anesthesia with thiopental 40mg/kg or isoflurane (4% for induction and 2 % for maintenance) did not influence the anxiolytic-like effects of diazepam in the EPM, namely: diazepam significantly raised the percentage of open arms entries and the percentage of time spent in open arms and had no effect in the number of enclosed arms compared to the control group **(Fig. 3).** The statistical data were detailed in table S4 of the supplementary material.

**Fig. 3.**
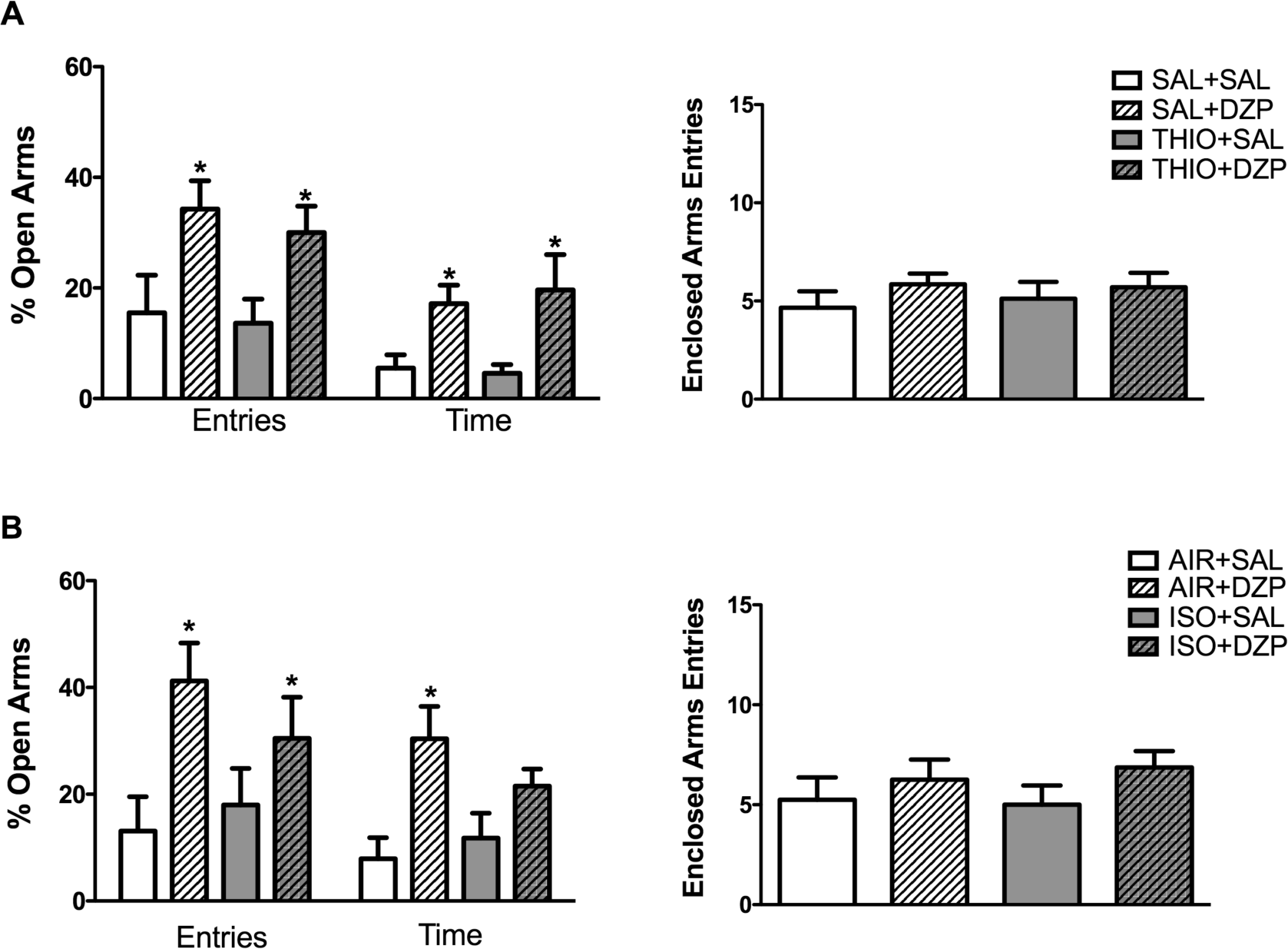
Previous anesthesia with thiopental or isoflurane does not affect the anxiolytic-like effect of diazepam in the elevated plus-maze (EPM). Anesthetics were injected 7 days and diazepam 1 h before the test. Rats were treated with (A) saline- saline (SAL-SAL, n=9); saline-diazepam 2.5mg/kg (SAL-DZP, n=7) thiopental 40mg/kg-saline (THIO-SAL, n=8); thiopental 40mg/kg-diazepam 2.5 mg/kg (THIO- DZP, n=10) and (B) air-saline (AIR-SAL, n=8); air-diazepam 2.5mg/kg (AIR-DZP, n=8); isoflurane-saline (ISO-SAL, n=8); isoflurane-diazepam 2.5mg/kg (ISO-DZP, n=8). *p<0,05 versus Sal-Sal group (two-way ANOVA followed by Duncan’s *post hoc* test).

In contrast, chloral hydrate 400mg/kg reduced the total distance moved in the OF 2 h after anesthetic administration compared to the control group. The other drugs did not modify the total distance traveled in the OF in none of the experiments (data and statistical analysis available in the tables S5, S6 and S7 of the supplementary material).

## 4 Discussion

This study aimed to evaluate whether 4 general anesthetics commonly used in procedures preceding psychopharmacological tests – tribromoethanol, chloral hydrate, thiopental and isoflurane - affect behavior in animal models predictive of antidepressant or anxiolytic-like effects. The main finding of our study was that previous anesthesia with the aforementioned anesthetics did not interfere with FST behaviors. To strengthen our findings, we evaluated whether previous anesthesia with these anesthetics would disturb the antidepressant-like effect of imipramine in rat FST paradimg. We showed that the antidepressant-like effect of imipramine in the FST was not influenced by previous anesthesia with tribromoethanol, chloral hydrate, thiopental or isoflurane. On the other hand, rats previously anesthetized with tribromoethanol or chloral hydrate exhibited, respectively, an anxiogenic-like or an anxiolytic-like profile in the EPM.

These results provide evidence that the choice of a specific anesthetic in situations that precede screening of anxiety-like effects should be careful. Moreover, as previous anesthesia with thiopental or isoflurane did not produce any *per se* effect in the EPM, we investigated their ability to obscure the anxiolytic-like effect of diazepam. The anxiolytic-like effect of diazepam was not affected by previous anesthesia with thiopental or isoflurane. Finally, as some studies suggest that low doses of specific general anesthetics produce antidepressant or anxiolytic-like effects (Gladney et al. 1994; Ozer et al. 2006; Engin et al. 2009), we included in our experimental design the investigation of the effects induced by subanesthetic doses.

### 4.1 Tribromoethanol effects

The general anesthetic tribromoethanol did not change rat behavior in the FST either 2 hours or 7 days after its administration. Despite this, we can rule out the possibility of a false negative, since imipramine, a tricyclic antidepressant, decreased immobility and increased climbing frequency in the FST. Also, previous anesthesia with tribromoethanol did not prevent the detection of the antidepressant-like effect of imipramine in the FST. Altogether, these results suggest that the use of tribromoethanol in surgeries preceding tests for screening of antidepressant-like effect of drugs is adequate.

Contrasting with data from FST, data from the EPM showed that previous anesthesia with tribromoethanol reduced open arms entries, suggesting a delayed anxiogenic-like effect. None of the tribromoethanol doses changed the number of enclosed arms entries in the EPM or the distance travelled in the OF, which means that there was no impairment of locomotor activity that could blunt the behaviors evaluated in the FST or in the EPM. Accordingly, under the tested conditions, tribromoethanol anesthesia seems not to be suitable in surgeries preceding behavioral tests that include the investigation of anxiety-related behaviors.

Besides that, a subanesthetic dose of tribromoethanol (90mg/kg) induced an anxiolytic-like effect similarly to that of diazepam in the EPM. Corroborating our findings, a low-dose of another primary alcohol, ethanol, increased exploration of the open arms in the EPM and of the light compartment in the light-dark transition test (Acevedo et al. 2014). This anxiolytic-like effect of tribromoethanol may be mediated by inhibitory currents through GABA_A_ receptors (Zimmerman et al. 1994; Krasowski and Harrison 2000; Thompson and Wafford 2001), similar to the mechanism of the anxiolytic effect of diazepam.

It is worth noting that a single anesthetic dose of tribromoethanol seems to generate safe and sufficient anesthesia for short-term procedures (Meyer and Fish, 2005; Ajadi et al., 2013; Gopalan et al., 2005; Pachon et al., 2015). However, its repeated use in rodents was associated with abdominal toxicity (Zeller et al., 1998; Maheras and Gow, 2013). Further, incorrect preparation or storage of tribromoethanol stocks increases the risk of mortality (Zeller et al. 1998; Weiss and Zimmermann 1999). Therefore, the rational for using this general anesthetic should take into account the different experimental circumstances in laboratory surgeries.

### 4.2 Chloral hydrate effects

Anesthesia with chloral hydrate did not change FST behaviors 2 hours or 7 days after its administration. In the same line, the antidepressant-like effect of imipramine was detected even in rats that received an anesthetic dose of chloral hydrate 7 days before the test. These results indicate that chloral hydrate may be safely administered in surgeries preceding behavioral analysis related to antidepressant-like effects.

The subanesthetic dose (150mg/kg) of chloral hydrate reduced the frequency of immobility 2 hours and 7 days after its administration. These effects raise the possibility of an acute and long-lasting antidepressant-like effect of a subanesthetic dose of chloral hydrate. Chloral hydrate did not significantly enhance any of the active behaviors (swimming and climbing), thus limiting the discussion about possible mechanisms of action related to its antidepressant effect. As evaluating the mechanism underlying this effect is beyond the scope of the present study, further studies are necessary.

Previous anesthesia of rats with chloral hydrate induced acute and long-lasting anxiolytic-like effect in the EPM. However, the animals that received the anesthetic dose of chloral hydrate presented a decrease in total distance traveled in the OF 2 hours after anesthetic administration, suggesting an impairment of locomotor activity. This impairment does not seem to interfere in the behaviors elicited by the EPM, since the exploration of enclosed arms, an index of EPM’s locomotor activity (Cruz et al. 1994), was similar between treated and control animals. Besides that, total distance traveled in the OF 7 days after chloral hydrate anesthesia was unchanged.

Other studies showed that a subanesthetic dose of this general anesthetic enhanced the number of transitions in the light-dark test and the punished crossings in four plates-test in mice (Aron et al. 1971; Gladney et al. 1994). The facilitative action of chloral hydrate in GABA_A_ receptors (Garrett and Gan 1998) could explain the possible anxiolytics effects observed in the EPM. Thereafter, our result suggests that chloral hydrate is not suitable for surgeries preceding behavioral tests related to anxiety.

In rodents, chloral hydrate seems to generate an effective and prolonged anesthesia (1 hour and 30 minutes approximately) (Field et al., 1993; Maud et al., 2014). Thus, the use of this anesthetic is advantageous for long-term procedures, because it may not be necessary to reapply the anesthetic during the procedure. On the other hand, some studies have described abdominal irritation and even abdominal necrosis by high doses of chloral hydrate (Dada et al. 1992; Vachon et al. 2000; Hüske et al. 2016). An additional issue relates to the poor analgesic effect of chloral hydrate (Flecknell, P 2009). Nevertheless, based on our data, the use of choral hydrate as an anaesthetic preceding behavioral studies should be avoided.

### 4.3 Thiopental effects

Anesthesia with thiopental acutely reduced the frequency of immobility and increased the frequency of swimming in the FST without altering distance traveled in the OF. In contrast, 7 days after anesthesia, FST exposed rats exhibited behaviors similar to the control group. As thiopental interferes with behavior in FST only acutely, this anesthetic might be administered in surgeries that precede behavioral tests for antidepressant-like effect screening. Reinforcing our statement, prior anesthesia with thiopental did not disturb the detection of the antidepressant-like effect of imipramine in rat FST.

Regarding the increase in the swimming frequency induced by thiopental, Detke and colleagues (Detke et al. 1995; Detke and Lucki 1996) suggested that enhancement of serotonergic transmission mediates swimming behavior. This hypothesis is based on the finding that selective serotonin reuptake inhibitors increase rat’s swimming frequency. Therefore, the antidepressant-like effect of thiopental may also be mediated by enhancement of serotonergic neurotransmission. In fact, at least in another mammal (*Arcicunthis niloticus*), anesthesia with thiopental augment total brain serotonin acutely (Mohamed Fathi et al. 1987). Also, thiopental appears to be a promiscuous injectable anesthetic not only potentiating GABA_A_ receptors function but also blocking nicotinic, 5-HT_3_ and NMDA receptors (Zhan et al. 1998; Downie et al. 2000). Given the complexity of thiopental actions, the specific mechanism of action underlying the antidepressant-like effect of thiopental needs to be investigated.

Unexpectedly, subanesthetic and anesthetic doses of thiopental did not change behaviors in the EPM, either acutely or 7 days after its administration. Moreover, previous anesthesia with this barbiturate seems not to interfere with the detection of the anxiolytic-like effect of diazepam in the EPM. Corroborating our results, pentobarbital, another barbiturate, also did not produce any change in rat’s behavior in the EPM (Luo et al. 2015). On the other hand, Torres et al. (1996) showed an acute anxiolytic-like effect of a subanesthetic dose of thiopental in rats tested in the one-way-avoidance test. Based on our results, thiopental appears to be a suitable anesthetic to be used in surgeries preceding behavioral tests for drugs that affect anxiety-related behaviors.

Thiopental is an ultra-short-acting anesthetic, therefore properly used for induction of general anesthesia (Flecknell, P 2009). On the other hand, during procedures such as rodent stereotaxic surgeries, thiopental commonly needs to be administered repeatedly, especially if it is used as a single anesthetic. Marked cumulative effects occur after repeated administration of this barbiturate, leading to a gradual prolongation of hypnosis with each successive injection and increasing risk of toxicity (Gepts and Camu 1991). The main adverse effects of thiopental are respiratory and cardiovascular depression (Kaczmarczyk and Reinhardt 1975; Suria et al. 1988; Gargiulo et al. 2012). Due to its restricted therapeutic index, thiopental use should be limited to situations in which other anesthetics are not recommended (Tagawa and Sakuraba 2014; Gonca 2015). Accordingly, although previous anesthesia with thiopental preceding behavioral evaluation related to depression and anxiety had proved to be suitable, this anesthetic should be chosen with caution.

### 4.4 Isoflurane effects

None tested concentration of isoflurane altered rat behavior in the FST, 2 hours or 7 days after anesthetic exposure. Yonezaki et al. (2015) also showed that 2 hours of isoflurane anesthesia do not induce any change in mice behavior evaluated in the FST and in the tail suspension test, one week after treatment. Conversely, a recent study showed an acute decrease in the immobility time in the FST induced by isoflurane anesthesia in mice (Antila et al. 2017). Although the anesthetic concentration was pretty much the same between the aforementioned studies, many other procedural differences may explain these conflicting results. While we tested rats 2 hours after isoflurane anesthesia that lasted 20 minutes, Antila and colleagues (2017) tested mice 15 minutes after isoflurane anesthesia that lasted 30 minutes. The same authors showed that previous (6 days before) anesthesia with isoflurane decreased failures to escape in the rat learned helpless paradigm, indicating a possible long-lasting antidepressant-like effect of isoflurane (Antila et al. 2017). Looking at all these results together, there appears to be a critical time window of exposure to isoflurane for the induction of the antidepressant-like effect. Given that exposure to the anesthetic isoflurane for a short period of time did not interfere in the detection of the antidepressant-like effect of imipramine in rats, under the conditions tested herein, previous anesthesia with isoflurane seems to be reliable in behavioral evaluations for screening antidepressant-like effect of drugs.

The subanesthetic and anesthetic concentrations of isoflurane had no effect in the EPM behaviors or in the locomotor activity acutely or after 7 days of exposure. In mice, anesthesia with isoflurane also did not interfere with behaviors in the EPM or in the light-dark test one week after anesthetic administration (Yonezaki et al. 2015). To strengthen our results, we evaluated whether prior anesthesia with isoflurane would disturb the anxiolytic-like effect of diazepam in the EPM. We showed that even in animals previously anesthetized with isoflurane, diazepam treatment was able to induce an anxiolytic-like effect. Consequently, isoflurane anesthesia proved to be viable in rats undergoing evaluation for anxiety-related effects.

Isoflurane is a volatile anesthetic that allows an appropriate control of anesthesia depth as well as rapid recovery. This anesthetic displays relatively safety for use in in both short and long-term procedures (Janssen et al. 2004; Cesarovic et al. 2010; Maud et al. 2014). Our study adds another advantage, which means, isoflurane appears to be an adequate anesthetic for use in surgeries preceding tests for antidepressant or anxiety-like effects. On the other hand, the needed of equipment for inhalant anesthetic administration enhances the cost and reduces convenience of anesthesia. Further, the waste anesthetic gas leakage may be a occupational risk for researchers (Todd et al. 2013; Johnstone et al. 2017). From these raised points, it is patent that several factors must be taken into account for the choice of general anesthetic in each experimental design.

### 4.5 Implications and limitations of the study

It is important to make some considerations about the experimental design of our study. We performed the behavioral tests on naive animals, which means that the animals were not exposed to surgical trauma. There are some evidences suggesting that surgical trauma impacts animal’s behavior (Li et al. 2010; Luo et al. 2015; Wang et al. 2017). Moreover, at least the effects of the anesthetic sevoflurane on anxiety might be affected by pain noxious stimulus (Luo et al. 2015). Our choice to evaluate the effects of the anesthetics in naïve animals was based on a strict control of anesthetic dose, given that during surgery, eventually, is necessary to reapply the anesthetic to ensure surgical anesthesia. Another reason to test naïve animals relates to an ethical concern. The administration of an anesthetic is mandatory in animals undergoing surgery, so that it would be impossible to have an adequate control group without anesthetic administration.

Another main limitation of our study refers to the fact that we investigated the effects of anesthesia merely in 2 behavioral tests. Despite this, the FST and the EPM present some features that make them appropriate choices for an initial screening. The FST paradigm has good predictive validity to detect antidepressant-like effect (Cryan et al. 2002). Further, changes in immobility time are consistent and easily reproduced in different laboratories, allowing to quickly performing a set of pharmacological experiments. Regarding the EPM paradigm, a valuable feature is its ability to detect both anxiogenic and anxiolytic-like effects of drugs (Kshama et al. 1990; Rodgers and Dalvi 1997; Griebel and Holmes 2013; Cipriano et al. 2016; Duarte et al. 2016). Given that anxiety is a heterogeneous emotional process and that the EPM seems to elicit unlearned anxiety, it would be interesting to expand the consequences of previous anesthesia to other paradigms of anxiety.

Finally, our study was restricted to evaluating the anesthetic effects in such specific conditions: naïve animals, single administration of selected anesthetics, fixed intervals of time between anesthesia and tests, evaluation of the effect of a single antidepressant or anxiolytic drug. In this line, our recommendations clearly are not valid to all behavioral protocols. Nevertheless, the present study reinforces the need for refinement of anesthetic’s choice.

## 5. Conclusions

Our results suggest that a single exposure to the general anesthetics thiopental or isoflurane seems to be suitable in surgeries that precede behavioral tests related to screening of antidepressant or anxiety-like effects. However, tribromoethanol and chloral hydrate are inadequate anesthetics for surgeries preceding behavioral tests involving anxiety evaluation due to its behavioral effects up to 7 days after its application. Finally, there appear to be no risk of bias in using tribromoethanol or chloral hydrate as anesthetics in surgeries prior behavioral tests that investigate antidepressant-like effect.

## Compliance with Ethical Standards

The authors declare that they have no conflict of interest. Ethical approval: All applicable international, national, and/or institutional guidelines for the care and use of animals were followed. Informed consent: Informed consent was obtained from all individual participants included in the study.

## Acknowledgements

LSH was recipient of a master student with research fellowship from Fundação de Amparo a Pesquisa e Inovaçao do Espirito Santo (FAPES) and TG was recipient of an undergraduate research fellowship from Universidade Federal do Espirito Santo (UFES). SJ receives a productivity fellowship from CNPq.

